# StrainFLAIR: Strain-level profiling of metagenomic samples using variation graphs

**DOI:** 10.1101/2021.02.12.430979

**Authors:** Kévin Da Silva, Nicolas Pons, Magali Berland, Florian Plaza Oñate, Mathieu Almeida, Pierre Peterlongo

## Abstract

Current studies are shifting from the use of single linear references to representation of multiple genomes organised in pangenome graphs or variation graphs. Meanwhile, in metagenomic samples, resolving strain-level abundances is a major step in microbiome studies, as associations between strain variants and phenotype are of great interest for diagnostic and therapeutic purposes.

We developed `StrainFLAIR` with the aim of showing the feasibility of using variation graphs for indexing highly similar genomic sequences up to the strain level, and for characterizing a set of unknown sequenced genomes by querying this graph.

On simulated data composed of mixtures of strains from the same bacterial species *Escherichia coli*, results show that `StrainFLAIR` was able to distinguish and estimate the abundances of close strains, as well as to highlight the presence of a new strain close to a referenced one and to estimate its abundance. On a real dataset composed of a mix of several bacterial species and several strains for the same species, results show that in a more complex configuration `StrainFLAIR` correctly estimates the abundance of each strain. Hence, results demonstrated how graph representation of multiple close genomes can be used as a reference to characterize a sample at the strain level.

**Availability:** http://github.com/kevsilva/StrainFLAIR

## INTRODUCTION

The use of reference genomes has shaped the way genomics studies are currently conducted. Reference genomes are particularly useful for reference guided genomic assembly, variant calling or mapping sequencing reads. For the later, they provide a unique coordinate system to locate variants, allowing to work on the same reference and easily share information. However, the usage of reference genomes represented as flat sequences reaches some limits (Ballouz et al., 2019).

Close reference genomes or genomes of strains from the same species show a high sequence similarity. Mapping sequencing reads on similar reference genomes results in mis-mapped reads or ambiguous alignments generating noise in the downstream analysis, that has yet to be clarified (Na et al., 2016). This has led recent methods to provide a representation of multiple genomes as genome graphs, also called variation graphs, in which each path is a different known variation. Such graph representations are well defined, and tools to build and manipulate graphs are under active development (Garrison et al., 2017; Kim et al., 2019; Rakocevic et al., 2019; Li et al., 2020).

This graph structure provides obvious advantages such as the reduction of the data redundancy, while highlighting variations (Garrison et al., 2018). However, it also introduces novel difficulties. Updating a graph with novel sequences, adapting existing efficient algorithms for read mapping, and, mainly, developing new ways to analyse sequence-to-graph mapping results for downstream analyses are among those new challenges. The work presented here primarily focuses on this latest point and proposes to show the feasibility of using a variation graph for identifying and estimating abundances, at the strain level, from an unknown metagenomic read set.

In the context of metagenomics, representing genomes in graphs is of particular interest for indexing microorganism genomes. Microorganisms are predominant in almost every ecosystems from ocean water (Sunagawa et al., 2015) to human body (Clemente et al., 2012), and play major functioning roles in them (New and Brito, 2020). While studies in microbial ecology are facing a bottleneck due to the difficulty of isolating and cultivating most of those microbes in laboratory, preventing the analysis of the complex structure and dynamics of the microbial communities (Stewart, 2012), high-throughput sequencing in metagenomics offers the opportunity to study a whole ecosystem. In particular, shotgun sequencing allows a resolution up to the species level (Jovel et al., 2016), and enable samples analysis in terms of population stratification, microbial diversity or bio-markers identification (Quince et al., 2017). Understanding of microbial communities structure and dynamics is usually revealed by resolving the species present in samples and their relative abundances, which can then be associated with phenotypes, notably in the field of human health (Ehrlich, 2011; Vieira-Silva et al., 2020; Solé et al., 2021). Now, characterizing samples at the strain level has a growing interest, as it may highlight new associations with phenotypes, and a better understanding of the functional impact of strains in host-microbe interactions is crucial to new therapeutic strategies and personalized medicine. *Escherichia coli*, which has a highly variable genome, is a well-known example since some strains are harmless commensals in the human gut microbiota while others are harmful pathogens (Rasko et al., 2008; Loman et al., 2013). Current approaches to handle multiple similar genomes as with strains use gene clustering and then select the representative sequence of each cluster, getting rid of the redundancy but also the variations, yet crucial to distinguish the strains of a species (Qin et al., 2010). Hence, indexation of a set of known strains is a good framework for testing the ability of a variation graph to capture the diversity while offering a way to correctly assign sequenced data to the strains they belong to.

In this work, we present `StrainFLAIR`, a novel method and its implementation that uses variation graph representation of gene sequences for strain identification and quantification. We proposed novel algorithmic and statistical solutions for managing ambiguous alignments and computing an adequate abundance metric at the graph node level. Results have shown that we could correctly identify and quantify strains present in a sample. Notably, we could also identify close strains not present in the reference.

`StrainFLAIR`is available at http://github.com/kevsilva/StrainFLAIR.

## METHODS

We propose here a description of our tool `StrainFLAIR` (STRAIN-level proFiLing using vArIation gRaph). This method exploits various state-of-the-art tools and proposes novel algorithmic solutions for indexing bacterial genomes at the strain-level. It also permits to query metagenomes for assessing and quantifying their content, in regards to the indexed genomes. An overview of the index and query pipelines are presented on Fig. 1.

**Figure 1.**
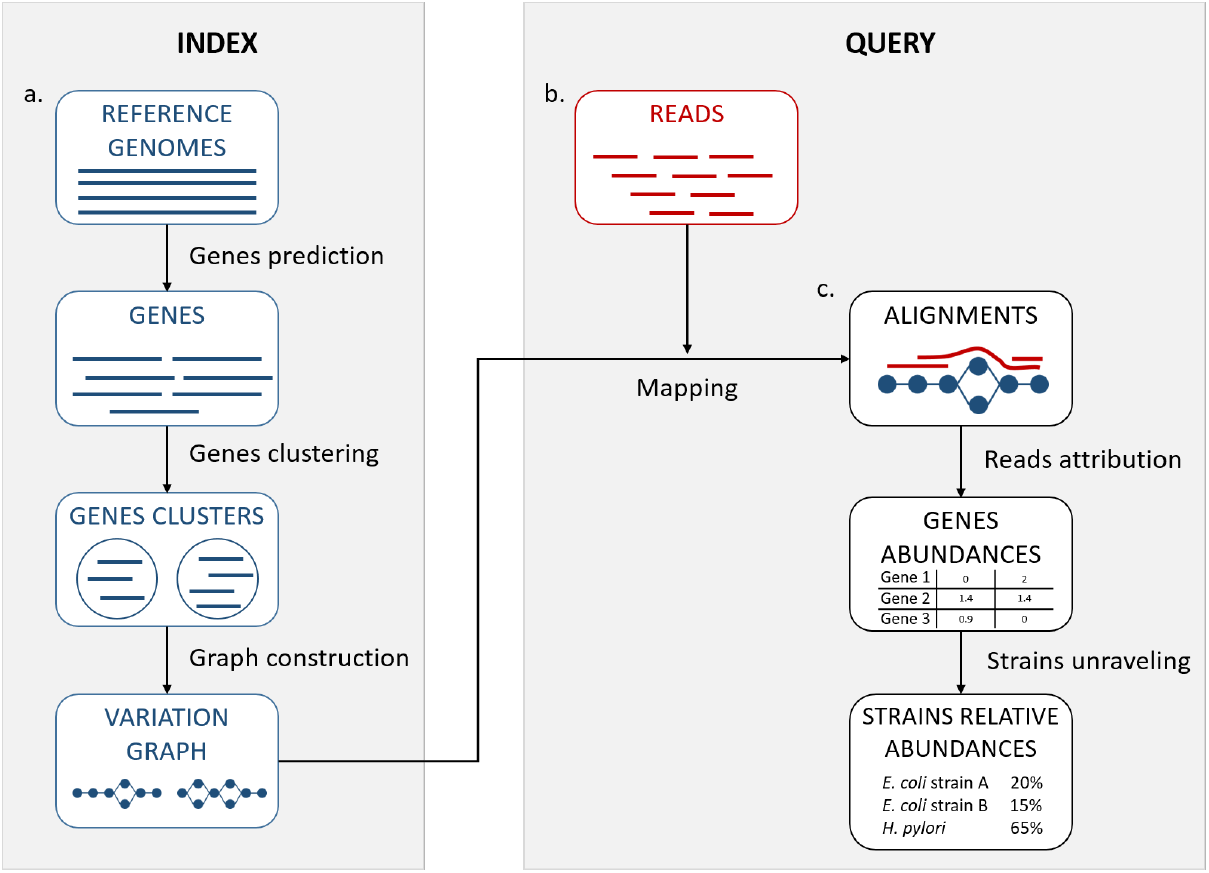
StrainFLAIR overview. a. Indexation. Input is a set of known reference genomes of various bacterial species and strains. `StrainFLAIR` uses a graph for indexing genes of those reference genomes. **b. Read mapping** on the previously mentioned graph. **c. Mapped reads analysis**. `StrainFLAIR` assigns and estimates species and strain abundances of a bacterial metagenomic sample represented as short reads.

Rational for the choice of third-party tools and their detailed usages are given in Supplementary Materials, Section S1.1.

### Indexing strains

#### Gene prediction

As non-coding DNA represents 15% in average of bacterial genomes and is not well characterized in terms of structure, `StrainFLAIR` focuses on protein-coding genes in order to characterize strains by their gene content and nucleotidic variations of them. Moreover, non-coding DNA regions can be highly variable (Thorpe et al., 2017) and taking into account complete genomes would then lead to highly complex graphs, and combinatorial explosions when mapping reads. Additionally, complete genomes are not always available. Focusing on the genes allows to use also drafts and metagenome-assembled genomes or a pre-existing set of known genes (Qin et al., 2010; Li et al., 2014). Hence, `StrainFLAIR` indexes genes instead of complete genomes in graphs.

Genes are predicted using `Prodigal`, a tool for prokaryotic protein-coding genes prediction (Hyatt et al., 2010).

Knowing that some reads map at the junction between the gene and intergenic regions, by conserving only gene sequences, mapping results are biased towards deletions and drastically lower the mapping score. In order to alleviate this situation, we extend the predicted gene sequences at both ends. Hence, `StrainFLAIR` conserves predicted genes plus their surrounding sequences. By default, and if the sequence is long enough, we conserve 75 bp on the left and on the right of each gene.

#### Gene clustering

Genes are clustered into gene families using `CD-HIT` (Li and Godzik, 2006). For the clustering step, the genes without extensions are used in order to strictly cluster according to the exact gene sequences and no parts of intergenic regions. `CD-HIT-EST` is used to realize the clustering with an identity threshold of 0.95 and a coverage of 0.90 on the shorter sequence. The local sequence identity is calculated as the number of identical bases in alignment divided by the length of the alignment. Sequences are assigned to the best fitting cluster verifying these requirements.

#### Graph construction

Each gene family is represented as a variation graph (Fig. 2). Variation graphs are bidirected DNA sequence graphs that represents multiple sequences, including their genetic variation. Each node of the graph contains sub-sequences of the input sequences, and successive nodes draw paths on the graph. Paths corresponding to reference sequences are specifically called “colored paths”. Each colored path corresponds to the original sequences of a gene in the cluster.

**Figure 2.**
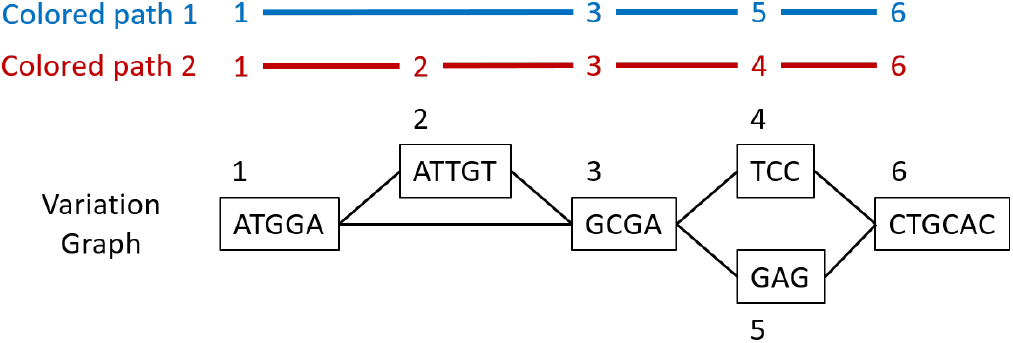
Illustration of a variation graph structure and colored paths. Each node of the graph contains a sub-sequence of the input sequences and is integer-indexed. A path corresponding to an input sequence is called a colored path, and is encoded by its succession of node ids, e.g. 1,3,5,6 for the colored path 1 in this example.

In the case of a cluster composed of only one sequence, `vg toolkit` (Garrison et al., 2017) is used to convert the sequence into a flat graph. Alternatively, when a cluster is composed of two sequences or more, `minimap2` (Li, 2018) is used to generate a multiple sequence alignment. Then `seqwish` (Garrison, 2021) is used to convert this multiple sequence alignment into a variation graph. All the so-computed graphs (one per input cluster) are then concatenated to produce a single variation graph where each cluster of genes is a connected component.

The index is created once for a set of reference genomes. Afterward, any set of sequenced reads can to be profiled at the strain-level based on this index.

### Querying variation graphs

#### Mapping reads

For mapping reads on the previously described reference graph, we use the sequence-to-graph mapper `vg mpmap` from `vg toolkit`. It produces a so-called “multipath alignments”. A multipath alignment is a graph of partial alignments and can be seen as a sub-graph (a subset of edges and vertices) of the whole variation graph (see Fig. 3 for an example). The mapping result describes, for each read, the nodes of the variation graph traversed by the alignment and the potential mismatches or indels between the read and the sequence of each traversed node.

**Figure 3.**
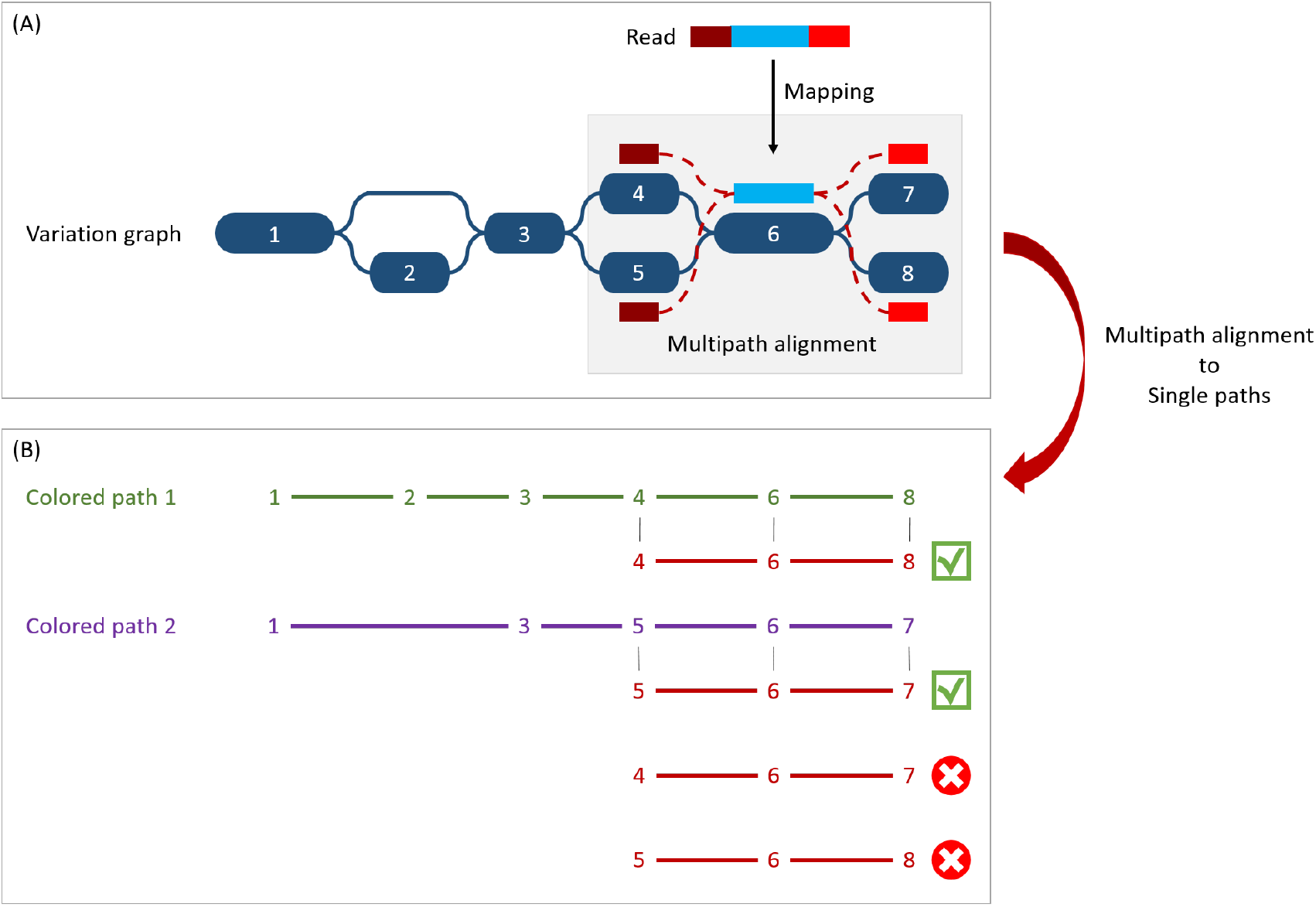
Illustration of the multipath alignment concept and the read attribution process. **(A) Path attribution**. The region of the read in blue aligns un-ambiguously to a node of the graph while the dark and light red parts can either align to the top or the bottom nodes of their respective mapping localization (due to mismatches that can align on both nodes for example), drawing an alignment as a sub-graph of the reference variation graph, and thus opening the possibility of four single path alignments. **(B) Colored path attribution**. First, from the multipath alignment (all four read sub-paths), the breadth search finds the possible corresponding single path alignments while respecting the mapping score threshold imposed by the user. Here, for the example, all four possible paths are considered valid. Second, each single path is compared to the colored paths from the reference variation graph. Two single path alignments matched the colored paths (4-6-8 and 5-6-7). As it mapped equally more than one colored path, this read falls in the multiple mapped reads case and is processed during the second step of the algorithm.

#### Reads attribution

When mapping a read on a graph with colored path, two key issues arise, as illustrated Fig. 3. As mapping generates a sub-graph per mapped read, the most probable mapped path(s) has / have to be defined. In the meanwhile, the most probable mapped path(s) corresponding to a colored path also have to be defined. Hence we developed an algorithm to analyse and convert, when possible, a mapping result into one or several continuous path(s) (successive nodes joined by only one edge) per mapped read. In addition we propose an algorithm to attribute such path to most probable colored path(s).

#### Path attribution

A breadth first search on the multipath alignment is proposed. It starts at each node of the alignment with a user-defined threshold on the mapping score. A single path alignment with a mapping score below this threshold is ignored, and the single path alignment with the best mapping score is retained. Additionally, for each alignment, nodes are associated with a so-called “horizontal coverage” value. The horizontal coverage of a node by a read corresponds to the proportion of bases of the node covered by the read. Hence, a node has an horizontal coverage of 1 if all its nucleotides are covered by the read with or without mismatches or indels.

Because of possible ties in mapping score, the search can result in multiple single path alignments, as illustrated Fig. 3(A). This situation corresponds to a read which sequence is found in several different genes or to a read mapping onto the similar region of different versions of a gene.

To take into account ambiguous mapping affectations, as shown below, the parsing of the mapping output is decomposed into two steps. The first step processes the reads that mapped only a unique colored path (called “unique mapped reads” here), corresponding to a single gene. The second step processes the reads with multiple alignments (called “multiple mapped reads” here).

#### Colored path attribution

Once a read is assigned to one or several path alignment(s), it still has to be attributed, if possible, to a colored path. The following process attributes each mapped read to a colored path and various metrics for downstream analyses are computed. In particular, an absolute abundance for each node of the variation graph, called the “node abundance”, is computed, first focusing on unique mapped reads (first step). For a given alignment, the successive nodes composing the path are compared to the existing colored paths of the variation graph. If the alignment matches part of a colored path, the number of mapped reads on this path is incremented by one (i.e. reads raw count). The node abundance for each node of the alignment is incremented with its horizontal node coverage defined by this alignment. Alignments with no matching colored paths are skipped.

Then, we focus on multiple mapped reads (second step), as illustrated Fig. 3(B). During this step, the alignment matches multiple colored paths. Hence, the abundance is distributed to each matching colored path relatively to the ratio between them. This ratio is determined from the reads raw count of each path from the first step. For example, if 70 unique mapped reads were found for path1 and 30 for path2 during the first step, a read matching ambiguously both path1 and path2 during the second step counts as 0.7 for path1 and 0.3 for path2. This ratio is applied to increment both the raw count of reads and the coverage of the nodes.

#### Gene-level and strain-level abundances

`StrainFLAIR` output is decomposed into an intermediate result describing the queried sample and gene-level abundances, and the final result describing the strain-level abundances.

#### Gene-level

After parsing the mapping result, the first output provides information for each colored path, *i*.*e*. each version of a gene. Thereby, this first result proposes gene-level information including abundances. Exhaustive description of these intermediate results is provided in Section S1.2 in Supplementary Materials. We describe here three major metrics outputted by `StrainFLAIR`:

The **mean abundance of the nodes composing the path**. Instead of solely counting reads, we make full use of the graph structure and we propose abundances computation for each node as previously explained, and as already done for haplotype resolution (Baaijens et al., 2019). Hence, for each colored path, the gene abundance is estimated by the mean of the nodes abundance. In order to not underestimate the abundance in case of a lack of sequencing depth (which could result in certain nodes not to be traversed by sequencing reads), the **mean abundance without the nodes of the path never covered by a read** is also outputted.

The mean abundance with and without these non-covered nodes are computed using unique mapped reads only or all mapped reads.

The **ratio of covered nodes**, defined as the proportion of nodes from the path which abundance is strictly greater than zero.

#### Strain-level

Strain-level abundances are then obtained by exploiting the specific genes of each reference genome from these intermediate results. First, for each genome, the proportion of detected genes is computed, as the proportion of specific genes on which at least one read maps. Then, the global abundance of the genome is computed as the mean or median of all its specific gene abundances. However, if the proportion of detected genes is less than a user-defined threshold, the genome is considered absent and hence its abundance is set to zero.

`StrainFLAIR` final output is a table where each line corresponds to one of the reference genomes, containing in columns the proportion of detected specific genes, and our proposed metrics to estimate their abundances (using mean or median, with or without never covered nodes as described for the gene-level result).

Results presented Section S1.3 in Supplementary Materials validate and motivate the proposed abundance metric by comparing it to the expected abundances and other estimations using linear models.

## RESULTS

We validated our method on both a simulated and a real dataset. All computations were performed using `StrainFLAIR`, version 0.0.1, with default parameters. The relative abundances estimation was based on the mean of the specific gene abundances, computed by taking into account all the nodes (including non-covered nodes), and using a threshold on the proportion of detected specific genes of 50%.

Results were compared to `Kraken2` (Wood et al., 2019) considered as one of the state-of-the-art tool dedicated to the characterization of read set content, and based on flat sequences as references. Read counts given by `Kraken2` were normalized by the genome length and converted into relative abundances.

Computing setup and performances are indicated in Supplementary Materials, Section S1.4.

### Validation on a simulated dataset

We first validated our method on simulated data, focusing on a single species with multiple strains. Our aim was to validate the `StrainFLAIR` ability to identify and quantify strains given sequencing data from a mixture of several strains of uneven abundances, and with one of them absent from the index.

#### Reference variation graph

We selected complete genomes of *Escherichia coli*, a predominant aerobic bacterium in the gut micro-biota (Tenaillon et al., 2010), and a species known for its phenotypic diversity (pathogenicity, antibiotics resistance) mostly resulting from its high genomic variability (Dobrindt, 2005).

Eight strains of *E. coli* were selected for this experiment from the NCBI^1^. Seven were used to construct a variation graph (*E. coli* IAI39, O104:H4 str. 2011C-3493, str. K-12 substr. MG1655, SE15, O157:H16 str. Santai, O157:H7 str. Sakai, O26 str. RM8426), and one was used as an unknown strain in a strains mixture (*E. coli* BL21-DE3).

#### Mixtures and sequencing simulations

Our aim was to simulate the co-presence of several *E. coli* strains. Two simulations with sequencing errors were conducted in order to highlight the detection and quantification of strains in a mixture. For each one, we tested our approach with various read coverage, as described below.

We simulated the sequencing of three strains to mimic complex single species composition in metagenomic samples. One of the strain was in equal abundance of one of the two others, potentially making it more difficult to distinguish, or in lower abundance, potentially making it more difficult to detect at all. The first simulation was a mixture composed of three strains contributing in the reference graph: *E. coli* O104:H4 2011c-3493, IAI39, and K-12 MG1655. The second simulation was a mixture composed of three strains: *E. coli* O104:H4 2011c-3493, IAI39, and BL21-DE3. The later being absent from the reference variation graph thus simulating a new strain to be identified and quantified.

For both simulations, short sequencing reads of 150 bp were simulated using `vg sim` from `vg toolkit` with a probability of errors set to 0.1%: 300,000 reads for *E. coli* O104:H4 2011c-3493 (representing ≈8.5x), 200,000 reads for *E. coli* IAI39 (representing ≈5.8x). For both simulations, various quantities of reads were generated for K-12 MG1655 or BL21-DE3: 200,000, 100,000, 50,000, 25,000, 10,000, 5,000 or 1,000 reads, representing approximately 6.5x, 3x, 1.6x, 0.8x, 0.3x, 0.2x, and 0.03x respectively for these two strains.

#### Strain-level abundances

As explained in Methods, we computed the strain-level abundances using the specific gene-level abundance table obtained by mapping the simulated reads onto the variation graph. We compared our results to the expected simulated relative abundances.

##### Simulation 1: mixtures with K-12 MG1655, present in the reference graph

`StrainFLAIR` successfully estimated the relative abundances of the three strains present in the mixture (Table 1), the sum of squared errors between the estimation given by our tool and the expected relative abundance was between 25 and 45 for all the experiments. However, it did not detect the very low abundant strain in the case of the mixture with 1,000 simulated reads for K-12 MG1655 (coverage of ≈ 0.03x). With our methodology, the threshold on the proportion of detected genes (see Methods) lead to set relative abundance to zero of likely absent strains. This reduces both the underestimation of the relative abundances of the present strains and the overestimation of the absent strains.

**Table 1.** Reference strains relative abundances expected and computed by StrainFLAIR or Kraken2 for each simulated experiment with variable coverage of the K-12 MG1655 strain. Best results are shown in bold. Complete results are presented Section S1.6 in Supplementary Materials.

| #reads<br>K-12 | Method | O104:H4 | IAI39 | K-12 | Sakai | SE15 | Santai | RM8426 |
| --- | --- | --- | --- | --- | --- | --- | --- | --- |
| 1,000 | Expected | 59.88 | 39.92 | 0.2 | 0 | 0 | 0 | 0 |
|  | StrainFLAIR | **56.45** | **43.55** | 0 | **0** | **0** | **0** | **0** |
|  | Kraken2 | 38.91 | 60.72 | **0.22** | 0.04 | 0.07 | 0.03 | 0.02 |
| 25,000 | Expected | 57.14 | 38.1 | 4.76 | 0 | 0 | 0 | 0 |
|  | StrainFLAIR | **52.1** | **40.58** | 7.32 | **0** | **0** | **0** | **0** |
|  | Kraken2 | 37.23 | 58.1 | **4.51** | 0.04 | 0.07 | 0.03 | 0.02 |
| 200,000 | Expected | 42.86 | 28.57 | 28.57 | 0 | 0 | 0 | 0 |
|  | StrainFLAIR | **38.12** | **29.83** | 32.05 | **0** | **0** | **0** | **0** |
|  | Kraken2 | 28.31 | 44.18 | **27.35** | 0.04 | 0.08 | 0.03 | 0.02 |

In comparison, `Kraken2` did not provide this resolution. Applied to our simulated mixtures, while `Kraken2` was slightly better for K-12 MG1655 abundance estimation, it overestimated IAI39 relative abundance and underestimated O104’s one, leading to an overall higher sum of squared errors (between 456 and 872) compared to the expected abundances. Moreover, it set relative abundances to all the seven reference strains whereas four of them were absent from the mixture. This was expected as some reads (from intergenic regions for example) can randomly be similar to regions of genes from absent strains.

##### Simulation 2: mixtures with BL21-DE3, absent from the reference graph

Here, BL21-DE3 was considered an unknown strain, not contributing to the variation graph. The closest strain of BL21-DE3 in the graph, according to fastANI (Jain et al., 2018), was K-12 MG1655 (98.9% of identity, see Supplementary Materials, Section S1.5). Thus we expected to find signal of BL21-DE3 through the results on K-12 MG1655.

As with the K-12 MG1655 mixtures, `StrainFLAIR` successfully estimated the relative abundances of the two known strains present in the mixture (Table 2), the sum of squared errors between the estimation given by our tool and the expected relative abundance was between 22 and 180 for all the experiments. Labelled as K-12, it also gave close estimations for BL21-DE3. Again, it did not detect the very low abundant strain in the case of the mixture with 1,000, 5,000, and 10,000 simulated reads for BL21-DE3. Also similarly to the K-12 MG1655 mixtures experiments, `Kraken2` overestimated IAI39 relative abundance and underestimated O104’s one (sum of squared errors between 751 and 873), even less

**Table 2.** Reference strain relative abundances expected and computed by StrainFLAIR or Kraken2 for each simulated experiment with variable coverage of the BL21-DE3 strain, absent from the reference variation graph. BL21-DE3 strain expected abundances are given in parentheses in the K-12 column. Best results are shown in bold. Complete results are presented Section S1.6 in Supplementary Materials.

| #reads<br>BL21-DE3 | Method | O104:H4 | IAI39 | K-12 | Sakai | SE15 | Santai | RM8426 |
| --- | --- | --- | --- | --- | --- | --- | --- | --- |
| 1,000 | Expected | 59.88 | 39.92 | (0.2) | 0 | 0 | 0 | 0 |
|  | StrainFLAIR | **56.47** | **43.53** | 0 | **0** | **0** | **0** | **0** |
|  | Kraken2 | 38.93 | 60.76 | **0.11** | 0.05 | 0.08 | 0.04 | 0.03 |
| 25,000 | Expected | 57.14 | 38.1 | (4.76) | 0 | 0 | 0 | 0 |
|  | StrainFLAIR | **54.09** | **41.71** | **4.2** | **0** | **0** | **0** | **0** |
|  | Kraken2 | 37.75 | 58.93 | 2.16 | 0.28 | 0.34 | 0.25 | 0.29 |
| 200,000 | Expected | 42.86 | 28.57 | (28.57) | 0 | 0 | 0 | 0 |
|  | StrainFLAIR | **46.95** | **35.34** | **17.72** | **0** | **0** | **0** | **0** |
|  | Kraken2 | 31.14 | 48.83 | 13.53 | 1.57 | 1.67 | 1.58 | 1.68 |

precisely than in the previous experiment. With sufficient coverage (here from the 0.8x for BL21-DE3), `StrainFLAIR` was closer to the expected values for all the reference strains than `Kraken2`.

Interestingly, the proportion of detected specific genes for each strain (Fig. 4) seems to highlight a pattern allowing to distinguish present strains, absent strains and likely new strains close to the reference in the graph. According to the experiments with enough coverage (from 25,000 simulated reads for BL21-DE3), three groups of proportions could be observed: proportion of almost 100% (O104:H4 and IAI39: strains present in the mixtures and in the reference graph), proportion under 30-35% (Sakai, SE15, Santai, and RM8426: strains absent from the mixtures), and an in-between proportion around 60-70% for K-12 MG1655 (closest strain to BL21-DE3).

**Figure 4.**
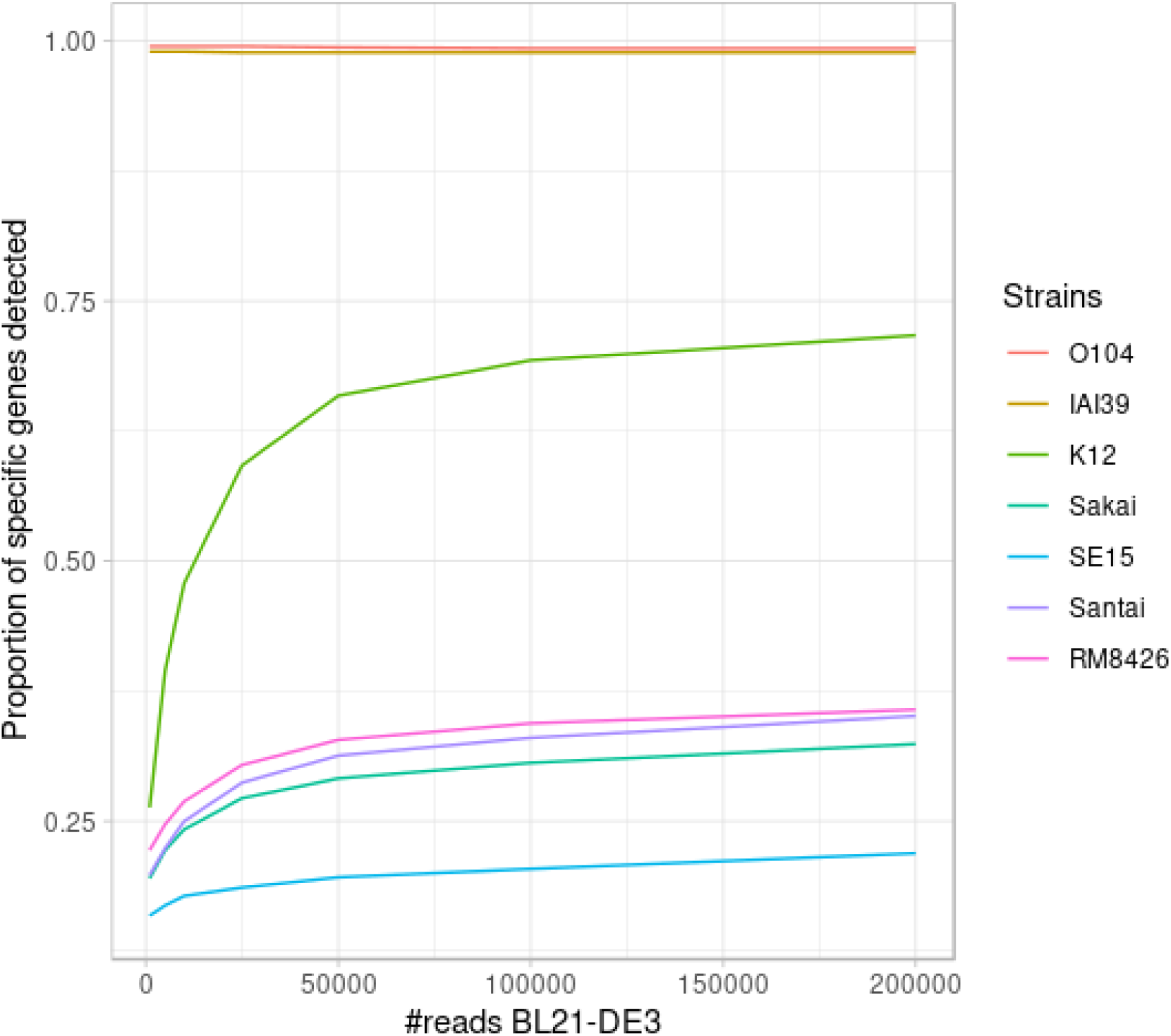
Proportion of detected specific genes for each simulated experiment with variable coverage of the BL21-DE3 strain, absent from the reference graph.

It was expected that an absent strain would have specific genes detected as `StrainFLAIR` detects a gene once only one read mappped on it. However, all absent strains had a proportion at around 30% except K-12 MG1655 which proportion was twice higher. Conjointly with the non-null abundance estimated for the reference K-12 MG1655, this suggests the presence of a new strain whose genome is highly similar to K-12 MG1655.

#### Validation on a real dataset

We used a mock dataset available on EBI-ENA repository under accession number PRJEB42498, in order to validate our method on real sequencing data from samples composed of various species and strains. The mock dataset is composed of 91 strains of bacterial species for which complete genomes or sets of contigs are available, including plasmids. Among the species, two of them contained each two different strains. Three mixes had been generated from the mock, and we used the “Mix1A” in the following results.

Even though 20 out of 91 strains were absents in this mix, we indexed the full set of 91 genomes. This was done in order to mimic a classical `StrainFLAIR` use case where the queried data is mainly unknown, and the reference graph contains species or strains not existing in these queried data. The metagenomic sample was sequenced using Illumina HiSeq 3000 technology and resulted in 21,389,196 short paired-end reads.

We compared our results to the expected abundances of each strain in the sample defined as the theoretical experimental DNA concentration proportion. As such, it has to be noted that potential contamination and/or experimental bias could have occurred and affected the expected abundances.

#### Strain detection

Among the 91 strains used in the reference variation graph, `StrainFLAIR` detected 65 strains. All of these 65 strains were indeed sequenced in Mix1A. Hence, `StrainFLAIR` produced no false positive. From the 26 strains considered absent by `StrainFLAIR`, 20 were not present in the sample (true negatives) and 6 should have been detected (false negatives). However, the term false negative has to be soften as the ground truth remains uncertain. Among those 6 undetected strains, all of them had theoretical abundance below 0.1%.

More precisely, among the 6 strains undetected by `StrainFLAIR`, 5 had some detected genes, but below the 50% threshold. In this case, by default, `StrainFLAIR` discards these strains. Finally, only one of the undetected strains (*Desulfovibrio desulfuricans* ND 132) should have been theoretically detected (even if its expected coverage was below 0.1%), but no specific gene was identified. Considering that `StrainFLAIR` uses a permissive definition of detected gene (at least one read maps on the gene), having strictly no specific genes detected for *Desulfovibrio desulfuricans* ND 132 suggests that this strain might in fact be absent from Mix1A. This is also supported by the result from `Kraken2` which estimated a relative abundance of ≈ 9e * 5, almost 500 times lower than the theoretical result.

As in the simulated dataset validation, `Kraken2` affected non-null abundances to all the references and thus could not be used to definitely conclude on presence/absence of strains in the sample.

#### Strain relative abundances

For the estimated relative abundances, `StrainFLAIR` gave more similar results compared to the state-of-the-art tool `Kraken2` than the experimental values (Fig. 5). The sum of squared error between `StrainFLAIR` and `Kraken2` was around 11. `StrainFLAIR` and `Kraken2` gave similar results compared to the experimental values, with sum of squared errors of around 209 and 211 respectively.

**Figure 5.**
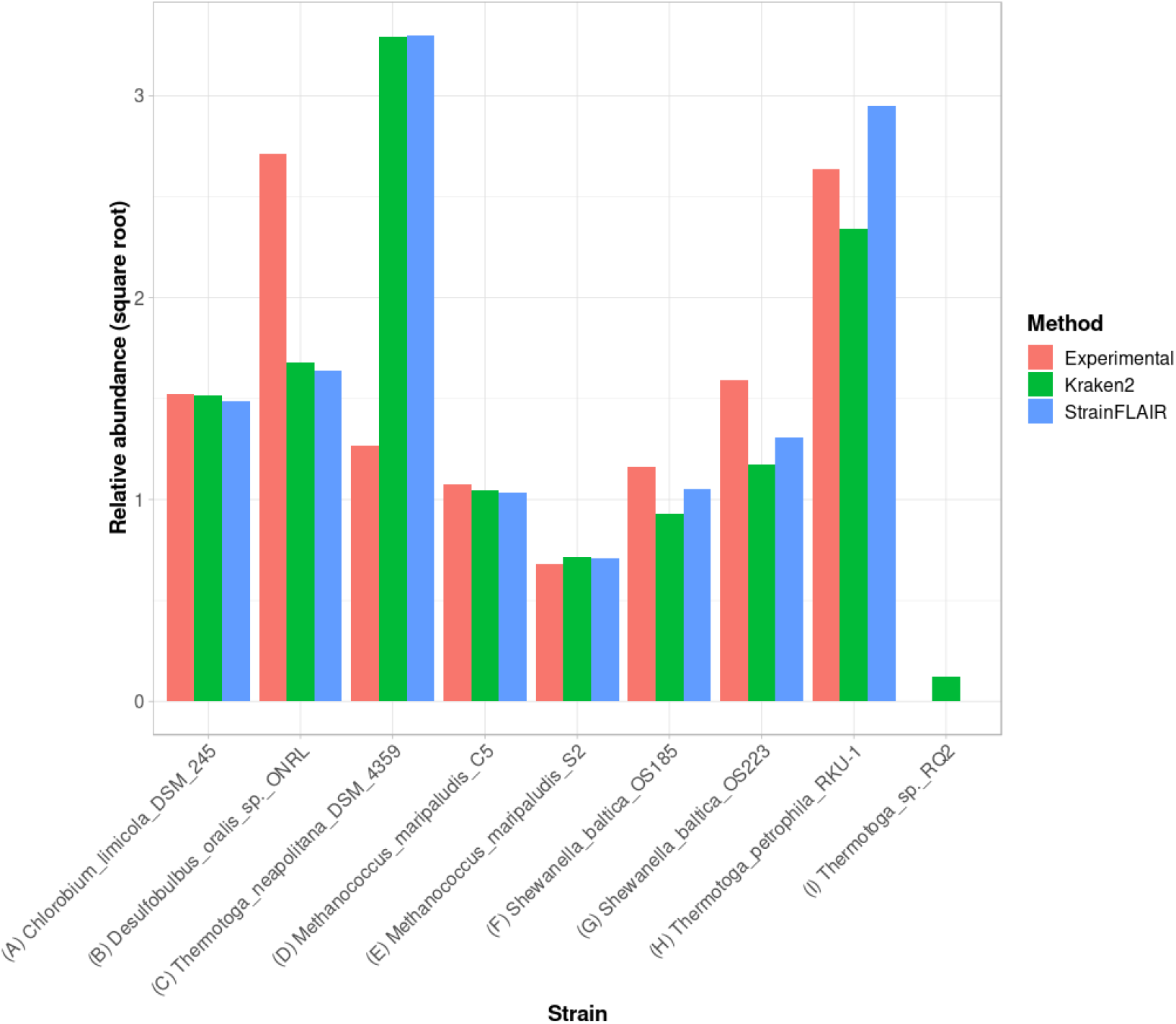
Experimental relative abundance compared to relative abundance as computed by StrainFLAIR and Kraken2. A selection of relevant results is shown here, see Supplementary Materials (Section S1.7) for the complete results. **(A)** Represents a case where `StrainFLAIR` and `Kraken2` give similar results to the experimental value (18 cases over 91). **(B)** Represents a case where `StrainFLAIR` and `Kraken2` give similar results, but lower than the experimental value (26 cases over 91). **(C)** Represents a case where `StrainFLAIR` and `Kraken2` give similar results, but greater than the experimental value (16 cases over 91). **(D, E, F, G)** Represent the two species represented by two strains each. **(H, I)** Represent two atypical cases.

Interestingly, *Thermotoga petrophila* RKU-1 is the only case where results from `StrainFLAIR` and `Kraken2` differs greatly, with, in addition, the theoretical abundance being in-between. Moreover, *Thermotoga* sp. RQ2 is the strain expected to be absent that `Kraken2` estimates with the highest relative abundance among the other expected absent strains, and the only one exceeding the relative abundances of two present strains. Considering the previous results on the simulated mixtures and that *Thermotoga petrophila* RKU-1 and *Thermotoga* sp. RQ2 are close species (fastANI around 96.6%) it could be an additional indicator of how tools like `Kraken2` can be mislead by too close species or strains.

In the sample, the species *Methanococcus maripaludis* was represented by two strains (S2 and C5) and the species *Shewanella baltica* likewise (OS223 and OS185). `StrainFLAIR` successfully distinguished and estimated the relative abundances of each strain of these two genomes. In this very situation and contrary to results on *E. coli* strains, `Kraken2` was also able to correctly estimate the abundances.

## DISCUSSION

Recent advances in sequencing technologies have provided large reference genome resources. Representation and integration of those multiple genomes, often highly similar, are under active development and led to genome graphs based tools. Integrating multiple genomes from the same species is particularly interesting as it provides new opportunities to characterize strains, a key resolution, for instance opening the field of precision medicine (Albanese and Donati, 2017; Marchesi et al., 2016).

In this context, we developed `StrainFLAIR`, a new computational approach for strain level profiling of metagenomic samples, using variation graphs for representing all reference genomes. Our intention was in the one hand to test whether or not indexing highly similar genomes in a graph enables to characterize queried samples at the strain level, and, in the other hand, to provide a end-user tool able to perform the indexation of genomes and the query of reads including the analyses of mapping results.

The method exploits state-of-the art-tools additionally to novel algorithmic and statistical solutions. By indexing microbial species and/or strains in a graph, it enables the identification and quantification of strains from a sequenced sample, mapped onto this graph.

We have demonstrated on simulated and on real datasets the ability of our method to identify and correctly estimate the abundance of microbial strains in metagenomic samples. In addition, `StrainFLAIR` was able to highlight the presence and also to estimate a relative abundance for a strain similar to existing references, but absent from these references.

We also showed that `StrainFLAIR` tended to set to zero the predicted abundance of low abundant strains, while a tool like `Kraken2` was able detect them. As a result, it seemed that `StrainFLAIR` looses the ability to detect very low abundant strains. However, in our simulations, this situation corresponded to coverages of 0.03x or less, hence simulating a strain for which not all genomic content was present. Eventually, it might be more relevant to define this strain as absent. Overall, there is a need to distinguish between low abundant strains, insufficient sequencing depth, and reads from intergenic regions or other genes randomly matching genes. In this regard, `StrainFLAIR` integrated a threshold on the proportion of specific genes detected that can be further explored to refine which strain abundances are set to zero. Importantly, results also showed that our graph-based tool had no false positive call, contrary to general purpose tool `Kraken2` that detected 100% of strains that were indexed but absent from queried reads.

From the validation on real datasets, we showed that `StrainFLAIR` was still able to correctly estimate the relative abundances in a more complex context mixing both different species and different strains, without being biased by references absent in the sample.

Our methodology taking into account all mapped reads and imposing a threshold that sets some strains abundances to zero seems more adequate and closer to what is expected in reality. Moreover, being able to detect some queried strains as absent is particularly interesting in the metagenomics context. Unlike mock datasets that are of controlled and known compositions, no prior knowledge is available for real metagenomic samples. They require the most exhaustive references - including unnecessary genomes - hence strains absent from the sample. `StrainFLAIR` is a new step towards the objective to take into account those unnecessary genomes without biasing the downstream analysis.

Measured computation time performances show that `StrainFLAIR` enables to analyse million reads in a few hours. Even if this opens the doors to routine analyses of small read sets, new development efforts will have to be made for reducing computation time in order to scale-up to very large datasets.

While `StrainFLAIR` focuses on profiling metagenomic samples at the strain level based on genes, it opens the way to pangenomic studies. Genome graphs are used to capture all the information on variation or similarity of sequences, which is particularly adapted to represent the gene repertoire diversity and the set of nucleotidic variations found between the different genomes of a species. This work highlights the importance to keep up working on pangenome graph representation.

The presence of queried unknown strain(s) is revealed both by reads mapping non-colored paths and by the amount of nucleotidic variations (indels and substitutions). The natural continuation will be related to the dynamical update of the graph when novel strains are detected in this way. This dynamicity will also be particularly useful considering the future flow of new sequenced metagenomes and the development of clinical metagenomics that will help to quickly and efficiently characterize in silico emerging strains of human health interest.

## ACKNOWLEDGMENTS

This work used the GenOuest bioinformatics core facility (https://www.genouest.org).

We acknowledge Mircea Podar for the providing of the mock dataset in premium access. Finally, we thank Mahendra Mariadassou, Rayan Chikhi, Olivier Jaillon and David Vallenet for all their advice along this work.

## S1 SUPPLEMENTARY MATERIALS

### S1.1 Third-party tools usage and rational

We propose here a the motivations and precise usage of the third-party tools that are employed in `StrainFLAIR`.

#### S1.1.1 Graph construction

`vg toolkit` allows to modify the graph including a normalization step. Normalization consists in deleting redundant nodes (nodes containing the same sub-sequence and having the same parent and child nodes), removing edges that do not introduce new paths, and merging nodes separated by only one edge.

For each cluster, if the colored paths of the corresponding graph still describe their respective input sequences, the graph is normalized.

After the concatenation of all computed graphs (one for each cluster), the final single variation graph is indexed using `vg toolkit`. Indexing a graph allows a fast querying of the graph when mapping reads. Indexation uses two file formats: *XG*, which is a succinct graph index which presents a static index of nodes, edges and paths of a variation graph, and *GCSA*, a generalized FM-index to directed acyclic graphs. A *SNARLS* file is also generated, describing snarls (a generalization of the superbubble concept (Paten et al., 2018)) in the variation graph and similarly allowing faster querying.

#### S1.1.2 Mapping reads

`vg toolkit` offers two sequence-to-graph mappers. The first one, `vg map`, outputs one or several final paths for each alignment. However, in case of several alignments with equal mapping scores, only one is randomly chosen. In order to get more exhaustive and accurate results, `StrainFLAIR` uses `vg mpmap` to map reads on the variation graph.

The mapping results are given in *GAMP* format, then converted into *JSON* format with `vg toolkit`, describing, for each read, the nodes of the graph traversed by the alignment.

### S1.2 Gene-level output by StrainFLAIR

Here we present the exhaustive description of information provided by `StrainFLAIR` at the gene level (before strain-level computations). For each colored path `StrainFLAIR` provides the following items:

- The corresponding gene identifier.
- For each reference genome, the number of copies of the gene. Since each unique version of a gene is represented once in the graph, whereas it can exist in several copies in the genome (duplicate genes), the counts and abundances computed correspond to the sum of those copies. Keeping track of the number of copies is important to normalize the counts.
- The cluster identifier to which the colored path belongs.
- For unique mapped reads: their raw number and their number normalized by the sequence length (see Section Querying variation graphs in Methods).
- For unique plus multiple mapped reads: their raw number and their number normalized by the sequence length (see Section Querying variation graphs in Methods).
- The mean abundance of the nodes composing the path, as defined in the manuscript.
- The mean abundance without the nodes of the path never covered by a read, as defined in the manuscript.
- The ratio of covered nodes, as defined in the manuscript.

### S1.3 Abundance metrics validation

The output of `StrainFLAIR` provides several metrics to estimate the abundance of the genes detected in the sample.

For validation, we used a combination of LASSO (least absolute shrinkage and selection operator) model and linear model on the simulated dataset to estimate the abundances at the strain-level, as the abundance of a gene is a linear combination of the abundances of the strains it belongs to. As such, Nwe expect no intercept value for those models and have forced the intercept at zero for the following modeling.

First, a LASSO model was used to perform strain selection. The response variable of the model was the presence or absence of the genes according to the selected metric while the strains, described as their genes content (number of copies), were the predictors. Then, a linear model was constructed with the raw selected metric as the response variable, and only the strains selected by the LASSO model as the predictors. The estimate of the strains relative abundance was thus the coefficients of the linear model associated to the strains and transformed into relative values. For each metric, the sum of squared errors between the real relative abundances and the estimated relative abundances from the linear model was computed. The best metric was then defined as the one minimizing this sum of squared errors.

For the mixtures containing *E. coli* K-12 MG1655, the three expected strains were selected and thus detected using LASSO, except for the mixture containing only 1,000 reads of K-12 MG1655 (representing 0.002% of the mixture, hence very negligible). For all the mixtures, the best metric was the mean abundance computed from the node abundances and by taking into account the multiple mapped reads.

For the mixtures containing *E. coli* BL21-DE3, BL21-DE3 being absent from the reference but very close to K-12 MG1655, we expected to get some detection of K-12 in the results. The three expected strains were selected and thus detected using LASSO, except for the mixture containing only 1,000 reads of BL21-DE3 (representing 0.002% of the mixture, hence very negligible). For the mixtures at 200,000, 100,000, and 50,000 reads of BL21-DE3, the best metric was the mean abundance computed from the node abundances without the abundances at zero, and by taking into account the multiple mapped reads. While for the others, the best metric was the mean abundance computed from the node abundances (including the abundances at zero), and by taking into account the multiple mapped reads.

This approach using linear models was particularly appropriate for this situation where the reference variation graph and the sample contained a small number of strains and thus a small number of predictors for the model. However, this can hardly transpose to a whole metagenomic sample with various species and various strains that would lead to too many predictors and probably confusing the heuristics behind the models. This was confirmed by applying the same methodology above on the mock dataset leading to abundances estimation hardly comparable to expected. Compared to `Kraken2` results, the sum of squared errors of our methodology was approximately 6 whereas for the results with the LASSO model it was around 236. Nevertheless, those results highlighted the relevance of (i) using a metric taking into account the multiple mapped reads and not only the unique mapped reads, and (ii) using our metric of abundance based on the node abundances over raw read counts.

### S1.4 Performances

Our benchmarks were performed on the GenOuest platform on a machine with 48 Xeon E5-2670 2.30 GHz with 500 GB of memory and 16 CPUs. Time results (Table S1) are the wall-clock times. We provided rough computation time, mainly in the purpose to show that `StrainFLAIR` can be applied on usual datasets.

**Table S1.**
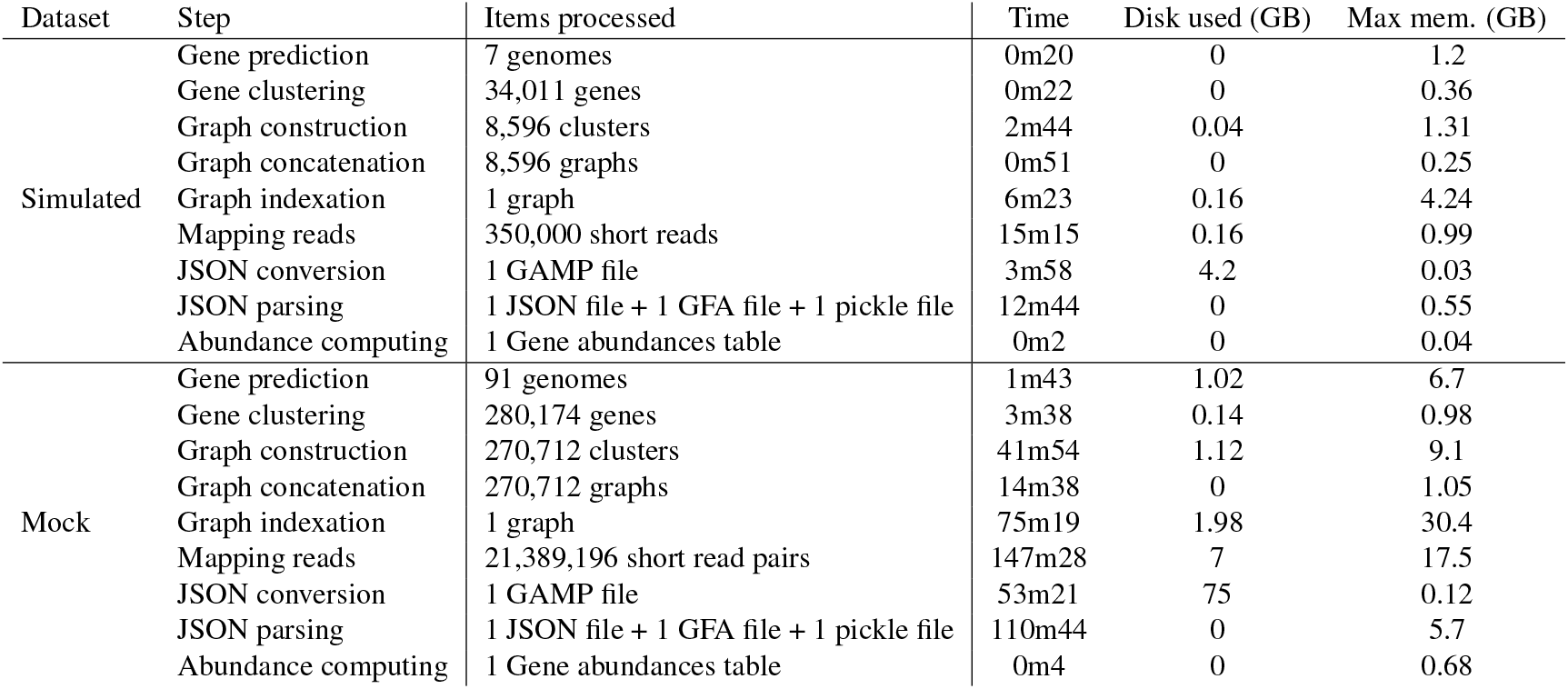
`StrainFLAIR` performances on simulated and mock datasets.

### S1.5 Distance between the selected genomes in the simulated experiment

We estimated the distance between the complete genomes of the selected strains using fastANI (Average Nucleotide Identity). FastANI uses an alignment-free algorithm to estimate the average nucleotide identity between pairs of sequences.

**Table S2.** Distance between each pair of complete genome sequences from eight strains of *E. coli* as computed by fastANI.

|  | K-12 | IAI39 | O104:H4 | Sakai | SE15 | Santai | BL21-DE3 | RM8426 |
| --- | --- | --- | --- | --- | --- | --- | --- | --- |
| K-12 | 100 | 97.0652 | 98.3769 | 97.8703 | 96.8716 | 98.0362 | 98.9365 | 98.3657 |
| IAI39 | 97.037 | 100 | 96.9742 | 96.7417 | 97.1289 | 96.9295 | 97.0197 | 96.8987 |
| O104:H4 | 98.3059 | 96.9521 | 100 | 97.4788 | 96.8007 | 97.8896 | 98.249 | 98.7212 |
| Sakai | 97.7497 | 96.8627 | 97.5094 | 100 | 96.6657 | 98.1523 | 97.7455 | 97.6125 |
| SE15 | 96.8453 | 97.1064 | 96.9211 | 96.7362 | 100 | 96.7575 | 96.8141 | 96.7763 |
| Santai | 98.0073 | 97.0372 | 97.9584 | 98.1797 | 96.8199 | 100 | 97.9279 | 97.9077 |
| BL21-DE3 | 98.9983 | 97.1721 | 98.4048 | 97.8227 | 96.8448 | 97.9616 | 100 | 98.3204 |
| RM8426 | 98.306 | 96.9037 | 98.6801 | 97.5815 | 96.6907 | 97.8353 | 98.2567 | 100 |

All pairs showed a distance at least greater than 95%, highlighting the strong similarities between the strains. As a threshold, we although considered that beyond 99%, sequences were too similar to be considered and distinguished, additionally to the effect of sequencing errors. The fastANI results showed that none of the pairs exceeded this similarity threshold.

The strain *E. coli* BL21-DE3 was chosen as the unknown strain while the seven others would be used to build the reference pangenome graph. According to the results of fastANI, the strain BL21-DE3 closest genome in the present references is the strain K-12 with a similarity of 98.9%. Hence we expected to find evidences of the strain K-12 while analyzing a sample containing the unknown strain BL21-DE3.

### S1.6 Detailed results from simulated datasets

**Table S3.** Reference strains relative abundances expected and computed by StrainFLAIR or Kraken2 for each simulated experiment with variable **coverage of the K-12 MG1655 strain**. Best results are shown in bold.

| #reads<br>K-12 | Method | O104:H4 | IAI39 | K-12 | Sakai | SE15 | Santai | RM8426 |
| --- | --- | --- | --- | --- | --- | --- | --- | --- |
| 1,000 | Expected | 59.88 | 39.92 | 0.2 | 0 | 0 | 0 | 0 |
|  | StrainFLAIR | **56.45** | **43.55** | 0 | **0** | **0** | **0** | **0** |
|  | Kraken2 | 38.91 | 60.72 | **0.22** | 0.04 | 0.07 | 0.03 | 0.02 |
| 5,000 | Expected | 59.41 | 39.6 | 0.99 | 0 | 0 | 0 | 0 |
|  | StrainFLAIR | **54.89** | **42.46** | 2.65 | **0** | **0** | **0** | **0** |
|  | Kraken2 | 38.61 | 60.25 | **0.99** | 0.04 | 0.07 | 0.03 | 0.02 |
| 10,000 | Expected | 58.82 | 39.22 | 1.96 | 0 | 0 | 0 | 0 |
|  | StrainFLAIR | **54.08** | **41.96** | 3.96 | **0** | **0** | **0** | **0** |
|  | Kraken2 | 38.26 | 59.69 | **1.9** | 0.04 | 0.07 | 0.03 | 0.02 |
| 25,000 | Expected | 57.14 | 38.1 | 4.76 | 0 | 0 | 0 | 0 |
|  | StrainFLAIR | **52.1** | **40.58** | 7.32 | **0** | **0** | **0** | **0** |
|  | Kraken2 | 37.23 | 58.1 | **4.51** | 0.04 | 0.07 | 0.03 | 0.02 |
| 50,000 | Expected | 54.55 | 36.36 | 9.09 | 0 | 0 | 0 | 0 |
|  | StrainFLAIR | **49.23** | **38.51** | 12.26 | **0** | **0** | **0** | **0** |
|  | Kraken2 | 35.63 | 55.6 | **8.62** | 0.04 | 0.07 | 0.03 | 0.02 |
| 100,000 | Expected | 50 | 33.33 | 16.67 | 0 | 0 | 0 | 0 |
|  | StrainFLAIR | **44.66** | **35.05** | 20.29 | **0** | **0** | **0** | **0** |
|  | Kraken2 | 32.8 | 51.19 | **15.85** | 0.04 | 0.07 | 0.03 | 0.02 |
| 200,000 | Expected | 42.86 | 28.57 | 28.57 | 0 | 0 | 0 | 0 |
|  | StrainFLAIR | **38.12** | **29.83** | 32.05 | **0** | **0** | **0** | **0** |
|  | Kraken2 | 28.31 | 44.18 | **27.35** | 0.04 | 0.08 | 0.03 | 0.02 |

Table S3 provides exhaustive results on simulated datasets when all queried strains are indexed in the variation graph. Table S4 provides exhaustive results on simulated datasets when one of the queried strain (BL21-DE3) is not indexed and highly similar to strain K-12.

**Table S4.** Reference strains relative abundances expected and computed by StrainFLAIR or Kraken2 for each simulated experiment with variable coverage of the BL21-DE3 strain, absent from the reference graph. BL21-DE3 being similar at 98.9% to K-12 strain (highest similarity compared to the other references), we expect that reads from BL21-DE3 will map this strain, hence its expected values are given in parentheses, as they correspond to BL21-DE3 strain abundances and not K-12. Best results are shown in bold.

| #reads<br>BL21-DE3 | Method | O104:H4 | IAI39 | K-12 | Sakai | SE15 | Santai | RM8426 |
| --- | --- | --- | --- | --- | --- | --- | --- | --- |
| 1,000 | Expected | 59.88 | 39.92 | (0.2) | 0 | 0 | 0 | 0 |
|  | StrainFLAIR | **56.47** | **43.53** | 0 | **0** | **0** | **0** | **0** |
|  | Kraken2 | 38.93 | 60.76 | **0.11** | 0.05 | 0.08 | 0.04 | 0.03 |
| 5,000 | Expected | 59.41 | 39.6 | (0.99) | 0 | 0 | 0 | 0 |
|  | StrainFLAIR | **56.45** | **43.55** | 0 | **0** | **0** | **0** | **0** |
|  | Kraken2 | 38.72 | 60.42 | **0.5** | 0.09 | 0.13 | 0.08 | 0.07 |
| 10,000 | Expected | 58.82 | 39.22 | (1.96) | 0 | 0 | 0 | 0 |
|  | StrainFLAIR | **56.45** | **43.55** | 0 | **0** | **0** | **0** | **0** |
|  | Kraken2 | 38.47 | 60.05 | **0.92** | 0.14 | 0.19 | 0.12 | 0.13 |
| 25,000 | Expected | 57.14 | 38.1 | (4.76) | 0 | 0 | 0 | 0 |
|  | StrainFLAIR | **54.09** | **41.71** | **4.2** | **0** | **0** | **0** | **0** |
|  | Kraken2 | 37.75 | 58.93 | 2.16 | 0.28 | 0.34 | 0.25 | 0.29 |
| 50,000 | Expected | 54.55 | 36.36 | (9.09) | 0 | 0 | 0 | 0 |
|  | StrainFLAIR | **52.74** | **40.62** | **6.65** | **0** | **0** | **0** | **0** |
|  | Kraken2 | 36.59 | 57.17 | 4.15 | 0.51 | 0.57 | 0.48 | 0.53 |
| 100,000 | Expected | 50 | 33.33 | (16.67) | 0 | 0 | 0 | 0 |
|  | StrainFLAIR | **50.47** | **38.64** | **10.89** | **0** | **0** | **0** | **0** |
|  | Kraken2 | 34.53 | 54.03 | 7.68 | 0.91 | 0.98 | 0.91 | 0.96 |
| 200,000 | Expected | 42.86 | 28.57 | (28.57) | 0 | 0 | 0 | 0 |
|  | StrainFLAIR | **46.95** | **35.34** | **17.72** | **0** | **0** | **0** | **0** |
|  | Kraken2 | 31.14 | 48.83 | 13.53 | 1.57 | 1.67 | 1.58 | 1.68 |

### S1.7 Detailed results for validation on mock datasets

**Figure S1.**
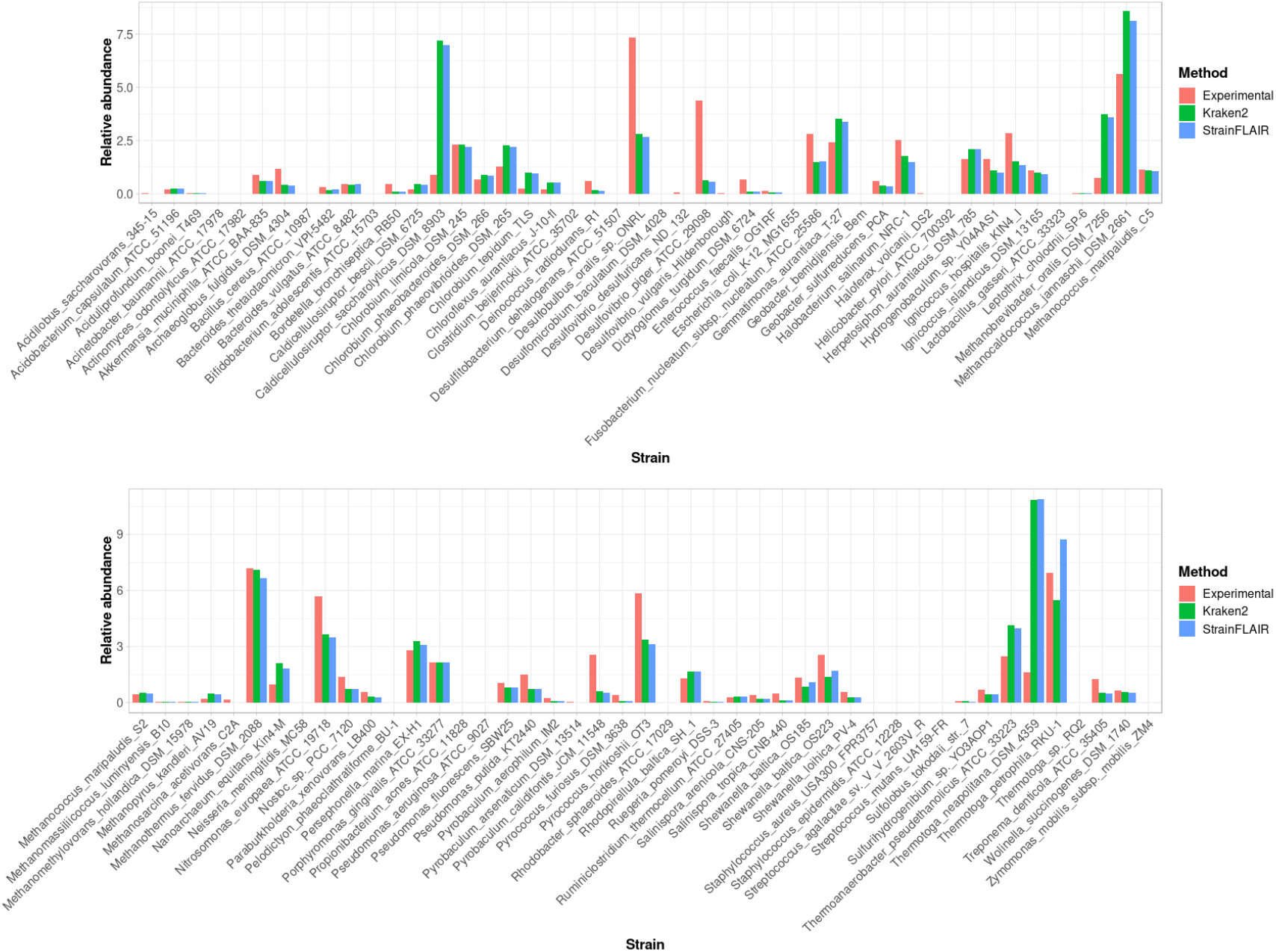
Experimental relative abundance compared to relative abundance computed by StrainFLAIR and Kraken2.

Figure S1 shows full results obtained on the mock dataset.

## Footnotes

1 https://www.ncbi.nlm.nih.gov/genome/?term=txid562[orgn]

